# Molecular characterisation of, and antimicrobial resistance in, *Clostridioides difficile* from Thailand, 2017-2018

**DOI:** 10.1101/2020.11.12.379040

**Authors:** Korakrit Imwattana, Papanin Putsathit, Daniel R Knight, Pattarachai Kiratisin, Thomas V Riley

**Affiliations:** School of Biomedical Sciences, The University of Western Australia, Western Australia; Faculty of Medicine Siriraj Hospital, Mahidol University, Thailand; School of Medical and Health Sciences, Edith Cowan University, Western Australia; School of Veterinary and Life Sciences, Murdoch University, Western Australia; Department of Microbiology, PathWest Laboratory Medicine, Queen Elizabeth II Medical Centre, Western Australia

**Author notes:** **Corresponding author -** Professor Thomas V Riley, School of Biomedical Sciences, The University of Western Australia, Nedlands, Western Australia, Australia.

## Abstract

**Background:** Antimicrobial resistance (AMR) plays an important role in the pathogenesis and spread of *Clostridioides difficile* infection (CDI). Many antimicrobials, such as fluoroquinolones, have been associated with outbreaks of CDI globally.

**Objectives:** This study aimed to characterise AMR among clinical *C. difficile* strains in Thailand, a country where the use of antimicrobials remains inadequately regulated.

**Methods:** Stool samples were screened for *tcdB* and positives were cultured. *C. difficile* isolates were characterised by toxin profiling and PCR ribotyping. Antimicrobial susceptibility testing was performed using an agar incorporation method, and whole-genome sequencing and AMR genotyping performed on a subset of strains.

**Results:** There were 321 *C. difficile* strains isolated from 326 stool samples. The most common toxigenic ribotype (RT) was RT 017 (18%), followed by RTs 014 (12%) and 020 (7%). There was a high resistance prevalence (≥ 10%) to clindamycin, erythromycin, moxifloxacin and rifaximin, and resistance prevalence was greatest among RT 017 strains. AMR genotyping revealed a strong correlation between resistance genotype and phenotype for moxifloxacin and rifampicin. The presence of *erm*-class genes was associated with high-level clindamycin and erythromycin resistance. Point substitutions on the penicillin-binding proteins (PBP1 and PBP3) were not sufficient to confer meropenem resistance, however, a Y721S substitution in PBP3 was associated with a slight increase in meropenem MIC. No resistance to metronidazole, vancomycin or fidaxomicin was observed.

**Conclusion:** There was a large proportion of *C. difficile* RT 017 in Thailand and a high AMR prevalence among these strains. The concordance between AMR phenotype and genotype was strong.

## Introduction

*Clostridioides* (*Clostridium*) *difficile* is a major cause of antimicrobial-associated diarrhoea globally (1). *C. difficile* infection (CDI) is a toxin-mediated disease and, so far, there have been three different major toxins identified: toxin A (TcdA), toxin B (TcdB) and binary toxin (*C. difficile* transferase, CDT). The genes encoding TcdA and TcdB are located on a 19.6 kb pathogenicity locus (PaLoc) (2) and the genes for CDT are located in a different part of the chromosome, the CDT locus (3). In non-toxigenic *C. difficile* (NTCD), the PaLoc is replaced by a fixed 115 bp locus (2). The toxin genes (*tcdA*, *tcdB*, *cdtA* and *cdtB*) can be detected by PCR (4, 5), however, some *C. difficile* strains, such as *C. difficile* ribotype (RT) 017, have a deletion at the repeating region of the *tcdA* gene, resulting in a truncated and non-functional toxin A (6). Several methods have been developed to detect this deletion (7, 8).

*C. difficile* can be separated into different RTs by amplifying the intergenic spacer region between the 16S and 23S rRNA genes (9). This classification method has been used worldwide due to its simplicity and high discriminating power (10). Important *C. difficile* RTs include *C. difficile* RT 027, an A+B+CDT+ strain associated with outbreaks of severe CDI in North America and Europe in the early 2000s (11), *C. difficile* RT 078, another A+B+CDT+ strain associated with the zoonotic transmission (12), and *C. difficile* RT 017, a *tcdA*-negative (A-B+CDT−) strain associated with global outbreaks since 1995 (6).

Although resistance to antimicrobials used for the treatment of CDI is relatively rare (13), resistance to other commonly used antimicrobials plays an important role in the pathogenesis of CDI and the spread of *C. difficile*. While intrinsic resistance to cephalosporins was probably responsible for an increase in the rate of CDI worldwide in the 1980s (14), resistance to clindamycin, new generation fluoroquinolones, rifampicin and tetracycline has been associated with outbreaks of CDI (15). These antimicrobials are also associated with an increased risk of developing CDI in general (16). Strict regulation of antimicrobials can be a successful measure to control CDI. In the US, such regulation has resulted in a significant decrease in CDI cases and CDI-related deaths over the last decade (17). Fluoroquinolone regulation in Australia has resulted in a relatively low prevalence of fluoroquinolone-resistant organisms (18), including *C. difficile* (19).

In previous studies, the epidemiology of CDI in Thailand has been characterised by a high prevalence of A-B+CDT− and NTCD, and an absence of A+B+CDT+ strains (20–22). *C. difficile* strains isolated in Thailand, especially *C. difficile* RT 017, were resistant to many antimicrobial groups, reflecting the use and misuse of these antimicrobials in the country (23). This study provides an update on the characterisation and antimicrobial susceptibility of *C. difficile* isolated from a tertiary hospital in Bangkok, Thailand.

## Materials and Methods

### Isolation and characterisation of *C. difficile*

This study was undertaken on 326 diarrhoeal stools samples collected from patients with a high index of suspicion of having CDI being treated at Siriraj Hospital, a large teaching hospital in Bangkok, Thailand, during 2017 – 2018. All stool samples were screened for the presence of the *tcdB* gene using the BD Max Cdiff assay (Becton Dickinson, US) as a part of routine investigations at Siriraj Hospital, and stools that were *tcdB* positive were then sent to a reference laboratory in Perth, Western Australia, for further investigation.

At the reference laboratory, stool samples were processed as previously described, including enrichment culture in cooked meat broth supplemented with gentamicin, cefoxitin and cycloserine (24). Characterisation of *C. difficile* was performed by PCR ribotyping and toxin gene profiling. PCR ribotyping was performed using the method described by Stubbs *et al* (9). The banding patterns were compared to a local database consisting of more than 80 internationally recognised RTs, including 15 reference RTs from the European Centre for Disease Prevention and Control. Patterns that did not match strains in the database were given an internal nomenclature. Detection of *tcdA* and *tcdB,* and the binary toxin genes (*cdtA* and *cdtB*), was performed using the methods described by Kato *et al* (7) and Stubbs *et al* (5), respectively. All NTCD isolates in this study were confirmed as such by the absence of the PaLoc using the method described by Braun *et al* (*lok* PCR) (2).

All stool samples were checked also for colonisation with multiple *C. difficile* strains. Briefly, DNA extraction was performed on all enrichment broths. DNA was then screened with either *tcdB* (7) or *lok* PCR (2), based on the toxin profile of the first *C. difficile* strain isolated from the specimen (2, 7). For example, a specimen previously positive for toxigenic *C. difficile* (TCD) was screened with *lok* PCR for NTCD and vice versa. All PCR-positive broths were re-cultured and the second *C. difficile* strain characterised as described above.

### Antimicrobial susceptibility testing

Antimicrobial susceptibility testing (AST) was performed using the agar incorporation method as described by the Clinical and Laboratory Standards Institute (CLSI) against nine antimicrobials listed in **Table 1** (25). *C. difficile* ATCC 700057, *Bacteroides fragilis* ATCC 25285, *Eubacterium lentum* ATCC 43055 and *B. thetaiotamicron* ATCC 29741 were included as control strains. Susceptibility results were interpreted using the minimal inhibitory concentration (MIC) breakpoints listed in **Table 1** (25–29). *C. difficile* strains that were resistant to at least three different antimicrobial classes were classified as multidrug-resistant (MDR). Resistance to clindamycin and erythromycin was considered as resistance to a single class (macrolide-lincosamide-streptogramin B; MLS_B_).

**Table 1.** List of antimicrobials, test ranges and susceptibility breakpoints used in this study.

| Antimicrobial | Test range (mg/l) | Breakpoint (mg/l) |  |  | Reference |
| --- | --- | --- | --- | --- | --- |
|  |  | S | I | R |  |
| Clindamycin | 0.008 – 256 | ≤ 2 | 4 | ≥ 8 | (25) |
| Erythromycin | 0.06 – 256 | - | - | > 8 | (26) |
| Moxifloxacin | 0.06 – 64 | ≤ 2 | 4 | ≥ 8 | (25) |
| Amoxicillin/clavulanate | 0.015 – 16 | ≤ 4 | 8 | ≥ 16 | (25) |
| Meropenem | 0.12 – 16 | ≤ 4 | 8 | ≥ 16 | (25) |
| Rifaximin | 0.008 – 64 | - | - | > 32 | (27) |
| Metronidazole | 0.008 – 4 | ≤ 2 | - | > 2 | (28) |
| Vancomycin | 0.03 – 4 | ≤ 2 | - | > 2 | (28) |
| Fidaxomicin | 0.002 – 2 | - | - | > 1 | (29) |

### Whole-genome sequencing, high-resolution typing and antimicrobial resistance characterisation

A subset of *C. difficile* strains (n = 37) was selected for whole-genome sequencing (WGS) to explore possible antimicrobial resistance (AMR) genotypes. Genomic DNA was extracted, sequenced on an Illumina HiSeq platform which generated 150 bp pair-end reads with a median coverage of 73X and characterised by multi-locus sequence typing (MLST) as previously described (30). Clade assignment of a newly defined sequence type (ST) was confirmed by comparing the average nucleotide identity (ANI) with *C. difficile* strains 630 (clade 1, GenBank accession AM180355) and R20291 (clade 2, GenBank accession FN545816) using FastANI (31). Known accessory AMR genes were identified by interrogating the read files with SRST2 version 0.2.0 against ARGannot database version 3 (32, 33). Draft annotated genomes were interrogated on Artemis version 17.0.1 to look for additional accessory genes (34). The genomes were interrogated also for the presence of known point substitutions associated with resistance to carbapenems (substitution in penicillin-binding proteins PBP1 and PBP3), fluoroquinolones (substitution in GyrA and GyrB subunit of the gyrase enzyme) and rifaximin (substitution in RpoB enzyme) (35, 36). The genotypes were then compared with phenotypic susceptibility data.

### Data availability

All sequence data generated in this study have been submitted to the European Nucleotide Archive under BioProject PRJEB40974, sample accessions ERS5247348 – ERS5247384. Details of the genomes are available in the **Supplementary Table S1**. The two newly characterised resistance determinants were submitted to the Nomenclature Center for MLS_B_ Genes (37), and the sequences were submitted to GenBank [accession numbers MW269959 (*erm*(52) gene) and MW269960 (*mef*(G) gene)]. The two genomes containing the prototypes of the genes were submitted to Genbank under BioProject PRJNA679085, accessions JADPMU000000000 (MAR225, carrying *erm*(52)) and JADPMT000000000 (MAR272, carrying *mef*(G)).

### Statistical analysis

All statistical analyses were performed using online tools by Social Science Statistics available at https://www.socscistatistics.com/. A p-value ≤ 0.05 was considered statistically significant.

### Human research ethics approval

This study was approved by the Human Research Ethics Committee of The University of Western Australia (reference file RA/4/20/4704) and the Siriraj Institutional Review Board (protocol number 061/2558 [EC1]).

## Results

### Characterisation of Thai *C. difficile*

A total of 296 *C. difficile* strains were initially isolated from the stool samples. Forty-four of the original 326 PCR positive stool samples were negative for TCD by culture; 30 contained no *C. difficile* while from 14 only NTCD was cultured. The enrichment broths for these samples were re-screened for TCD and were all negative by *tcdB* PCR. Another 25 strains were identified from the co-colonisation screening process, yielding a total of 321 *C. difficile* strains. Of these, 221 (68.85%) were positive for *tcdA* and *tcdB* (A+B+CDT−), 58 (18.07%) were positive for *tcdB* only and had a deletion in *tcdA* (A-B+CDT−), three (0.93%) were positive for *tcdA*, *tcdB*, as well as *cdtA* and *cdtB* (A+B+CDT+) and 39 strains (12.15%) were negative for all toxin genes (A-B−CDT−, NTCD).

The 321 *C. difficile* strains belonged to 63 different RTs, 19 of which were internationally recognised. The remaining RTs were given internal nomenclature beginning with either “QX” or “KI”. The prevalence of the most common RTs is summarised in **Table 2**. The most common TCD strain was *C. difficile* RT 017 (A-B+CDT−), followed by RTs 014 and 020 (both A+B+CDT−). The most common NTCD was *C. difficile* RT 010.

**Table 2.** Ribotypes of 321 *C. difficile* strains from Thailand, by toxin profile.

| Toxin Profile | Ribotype | Number | % of toxigenic strains | % of all |
| --- | --- | --- | --- | --- |
| **A+B+CDT-** | **Total** | **221** | **78%** | **69%** |
|  | 014 | 40 | 14% | 12% |
|  | 020 | 24 | 9% | 7% |
|  | 046 | 18 | 6% | 6% |
|  | QX517 | 17 | 6% | 5% |
|  | 297 | 13 | 5% | 4% |
|  | QX026 | 11 | 4% | 3% |
|  | 043 | 10 | 4% | 3% |
|  | Others | 88 | 31% | 27% |
| **A-B+CDT-** | **Total** | **58** | **21%** | **18%** |
|  | 017 | 58 | 21% | 18% |
| **A+B+CDT+** | **Total** | **3** | **1%** | **1%** |
|  | 078 | 1 | <1% | <1% |
|  | QX273 | 1 | <1% | <1% |
|  | KI008 | 1 | <1% | <1% |
| **A-B-CDT-** | **Total** | **39** | **-** | **12%** |
|  | 010 | 7 | - | 2% |
|  | QX002 | 5 | - | 2% |
|  | 009 | 4 | - | 1% |
|  | QX011 | 4 | - | 1% |
|  | Others | 19 | - | 6% |

Note: RTs designated with "QX" and "KI" were RTs that did not match the internationally recognised RTs in the database and were given internal nomenclature.

### Characterisation of a novel binary toxin-positive *C. difficile* isolate

One of the *C. difficile* strains isolated in this study was positive for all three toxin genes (A+B+CDT+) and had a unique ribotyping pattern compared to the local reference library. According to the MLST scheme, this isolate was characterised as the novel ST 692 within evolutionary clade 1. However, pairwise ANI analysis showed that this strain was more closely related to *C. difficile* R20291 (clade 2, ANI = 99.17%) than *C. difficile* 630 (clade 1, ANI = 98.89%).

### Antimicrobial susceptibility of Thai *C. difficile*

Overall antimicrobial susceptibility results are shown in **Table 3** and the MIC distribution of selected six antimicrobial classes is displayed in **Figure 1**. Based on the MIC value, clindamycin-resistant *C. difficile* strains could be divided into two groups: those with MIC ≥ 32 mg/l (n = 97) and those with MIC < 32 mg/l (n = 166). There was a strong correlation between high-level clindamycin resistance and erythromycin resistance: 95 strains (97.94%) that had clindamycin MIC ≥ 32 mg/l were also resistant to erythromycin while only 16 strains (9.64%) in the other group were resistant to erythromycin (Cohen’s kappa = 0.857). There was also a clear separation in MIC value between strains with and without rifaximin resistance. The separation was less clear for moxifloxacin and was not observed in amoxicillin/clavulanate and meropenem.

**Table 3.** Antimicrobial susceptibility of 321 *C. difficile* strains from Thailand.

| Antimicrobial | MIC range (mg/l) | MIC<sub>50</sub> (mg/l) | MIC<sub>90</sub> (mg/l) | Susceptibility (%) |  |  |
| --- | --- | --- | --- | --- | --- | --- |
|  |  |  |  | S | I | R |
| Clindamycin | 0.5 – >256 | 8 | >256 | 26 (8%) | 32 (10%) | 263 (82%) |
| Erythromycin | 0.5 – >256 | 2 | >256 | - | - | 109 (34%) |
| Moxifloxacin | 0.5 – >64 | 2 | 32 | 242 (75%) | 3 (1%) | 76 (24%) |
| Amoxicillin/clavulanate | 0.25 – 8 | 0.5 | 1 | 320 (>99%) | 1 (<1%) | 0 |
| Meropenem | 1 – 16 | 4 | 8 | 280 (87%) | 39 (12%) | 2 (1%) |
| Rifaximin | ≤0.008 – >64 | 0.03 | 2 | - | - | 31 (10%) |
| Metronidazole | 0.06 – 1 | 0.25 | 0.25 | 321 (100%) | - | 0 |
| Vancomycin | 0.5 – 2 | 1 | 2 | 321 (100%) | - | 0 |
| Fidaxomicin | 0.03 – 0.25 | 0.125 | 0.25 | - | - | 0 |

**Figure 1.**
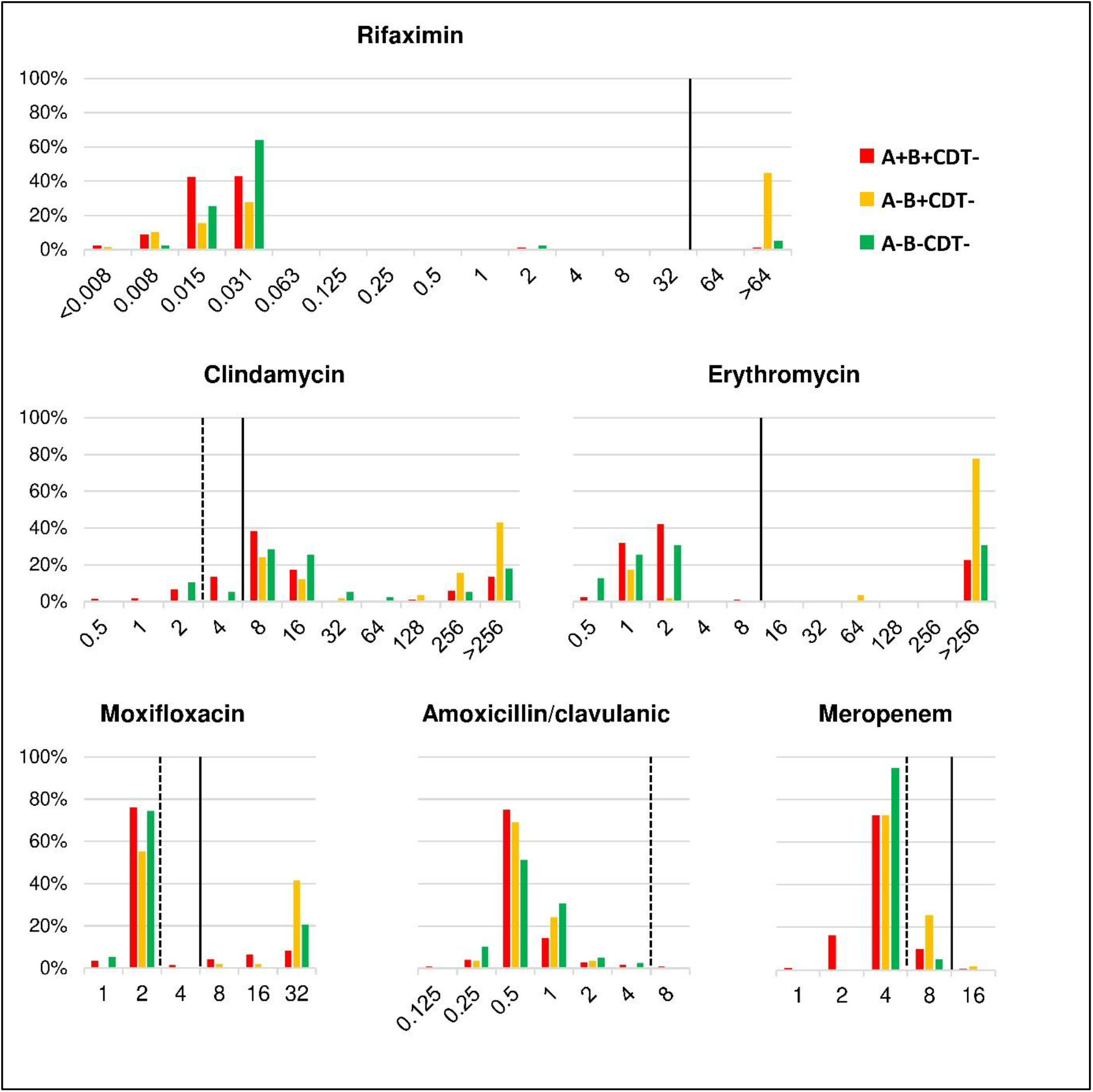
MIC distribution for six antimicrobials against 321 strains of C. difficile in Thailand. *C. difficile* strains were classified according to their toxin gene profiles: A+B+CDT−, red; A-B+CDT−, yellow; A-B−CDT−, green. The number of A+B+CDT+ C. difficile strains was low (n = 3) and these were excluded. Breakpoints for intermediate resistance (I) and resistance (R) are shown by broken and solid lines, respectively.

When classified by toxin gene profiles, resistance to clindamycin, erythromycin, moxifloxacin and rifaximin were more prevalent among A-B+CDT− *C. difficile*, all of which belonged to RT 017, than A+B+CDT- and NTCD (**Figure 1**). A total of 29 (9.03%) *C. difficile* isolates were MDR, 26 (8.10%) of which were *C. difficile* RT 017. The remaining isolates were NTCD (n=2) and A+B+CDT− *C. difficile* (n=1). All MDR isolates were resistant to the MLS_B_ group (both clindamycin and erythromycin), moxifloxacin and rifaximin. One MDR isolate was also resistant to meropenem (A-B+CDI−, RT 017, MIC = 16 mg/l).

### Genotypic resistance determinants in Thai *C. difficile*

A summary of MIC values and genotypic resistance determinants of 37 sequenced *C. difficile* strains is available in **Supplementary Table S2**. Of these strains, 31 had high-level resistance to clindamycin: 23 strains carried an *erm*(B) gene, five carried an *erm*(G) gene and three carried a gene encoding an rRNA adenine N(6)-methyltransferase family protein, the closest match to which in the CARD database was the Erm(42) protein [39% identity, E-value = 1.19E-53] (38). This newly characterised gene was given the name *erm*(52). Of the 23 *erm*(B)-positive strains, 19 carried the *erm*(B) gene on transposon Tn*6194* (82.61%), while the other four (17.39%) carried the gene on Tn*6189*. No *erm*-class genes were identified among strains with low-level clindamycin resistance. The concordance between the presence of *erm*-class genes and high-level clindamycin resistance was 100%. A gene encoding a macrolide efflux protein was identified in two strains with high-level erythromycin resistance (MIC > 256 mg/l), and given the name *mef*(G). No significant genotypic resistance determinants were identified in strains with low-level clindamycin resistance.

A T82I substitution in GyrA was found in 23 strains and a D426V substitution in GyrB found in one strain with a moxifloxacin MIC of 16 – 32 mg/l. There were no known point substitutions in the remaining 14 strains, 13 of which were moxifloxacin-susceptible; one had low-level moxifloxacin resistance (MIC 8 mg/l) [97.37% concordance]. There were H502N and R505K substitutions in RpoB in all 23 rifaximin-resistant strains and none of the susceptible strains [100% concordance].

Twelve strains had an A555T substitution in penicillin-binding protein 1 (PBP1) and another seven had a Y721S substitution in PBP3. A multiple linear regression analysis was performed to assess the association between the presence of these substitutions and the MIC values for meropenem. The Y721S substitution in PBP3 was associated with an increase in meropenem MIC (adjusted R^2^ = 0.516, t = 5.521, p < 0.0001), while the A555T substitution in PBP1 was not associated with the change in meropenem MIC (t = −1.127, p = 0.268).

## Discussion

This study provides an update on the molecular epidemiology and antimicrobial susceptibility of *C. difficile* strains circulating in Thailand. It also explores the genomic basis of important AMR in these strains. As the focus was on patients suspected of having CDI and not on the prevalence of *C. difficile* in the general population, the stool samples included in this study were first screened for the *tcdB* gene by PCR before culture. Thus the overall prevalence of TCD was higher than previous studies in Thailand (20–22), however, the common strains found were similar. The majority of A+B+CDT− strains belonged to *C. difficile* RT 014/020 group, all A-B+CDT− strains belonged to *C. difficile* RT 017 and most NTCD belonged to *C. difficile* RTs 009, 010 and 039. Three binary toxin-positive isolates were found in this study, one of which was *C. difficile* RT 078. The epidemic *C. difficile* RT 027 remained absent in Thailand despite its successful spread in other regions (39).

Why *C. difficile* RT 027 has failed to spread and to establish in Thailand, and Southeast Asia in general, remains unknown. One possible reason is that the successful spread of this RT was mainly due to its resistance to fluoroquinolones which provided a selective advantage over other less resistant RTs (40). Although there is high consumption of fluoroquinolones, such as levofloxacin, in the country (41), Thailand already harbours *C. difficile* RT 017, another epidemic RT many of which are resistant to fluoroquinolones, as well as other antimicrobials (15). Thus, it may have been difficult for *C. difficile* RT 027 to compete with this local RT compared to other regions.

Though *C. difficile* RT 027 was not identified in this study, a possible relative of this hypervirulent strain, ST 692, was isolated. The MLST for this strain was unusual, as it was classified into clade 1 despite containing binary toxin genes in a complete CDT locus. The presence of binary toxin genes is a feature common in *C. difficile* clades 2 and 5, but rare in clade 1 (42). Thus, an ANI analysis was performed, which indicated that this newly characterised strain was more related to clade 2 than clade 1 *C. difficile*, as expected from the toxin gene profile. Clades 1 and 2 *C. difficile* are closely related and share a large proportion of housekeeping gene alleles used in the MLST scheme. As a result, it may be difficult to properly discriminate these two clades by this method. The use of ANI analysis, which involves the whole genome rather than a specific set of housekeeping genes, can help in the correct classification of some borderline strains as shown in a previous study (42). According to the ANI analysis, it is more likely that this newly described *C. difficile* ST 692 belongs to clade 2 and is related to *C. difficile* RT 027.

A discordance between culture results and the result of a conventional real-time *tcdB* PCR was observed in 44 stool samples. The false-positive rate of the real-time PCR method (13.50%) was comparable to the previous report comparing *tcdB* PCR with a similar enrichment culture method but without the colonisation screening step (24). This suggests that the additional screening step does not significantly increase the yield of the culture method, although it may help identify stool samples with multiple *C. difficile* strains. This false-positive rate also highlights the importance of patient clinical data or additional tests to improve the accuracy of CDI diagnosis. In the latest guidelines for the treatment and diagnosis of CDI, *tcdB* PCR is no longer recommended as a stand-alone test unless the patient has symptoms suggestive of CDI (43).

AMR in *C. difficile* affects both the pathogenesis and treatment of CDI. To cause the disease, *C. difficile* must tolerate the presence of antimicrobials in the intestinal lumen while the microbiota perishes (15). Many successful *C. difficile* lineages have indeed been characterised with increased resistance to at least one major drug group (15). In this study, *C. difficile* RT 017, which was the most common RT, had greater resistance to MLS_B_ (both clindamycin and erythromycin), moxifloxacin and rifaximin than other RTs. It was also the most common MDR *C. difficile* strain in this study. *C. difficile* RT 017 has been reported also to be the most prevalent RT with significant resistance to many antimicrobials in other parts of Thailand (22). This particular RT has been associated with resistance to at least six antimicrobial groups (15), which may have accounted for its successful global spread (6). As regulation of antimicrobial use has reduced the impact of *C. difficile* in many countries (17, 19), a similar approach should be effective for the control CDI in Thailand.

All *erm*(B)-positive *C. difficile* strains carried the gene on two well-characterised *erm*(B)-positive transposons: Tn*6189* and Tn*6194*, the latter being found also in *C. difficile* M68, a *C. difficile* RT 017 strain widely used as a reference in genomic studies (15). Tn*6194*, the most prevalent transposon in this study, is capable of inter-species transfer, most notably between *C. difficile* and *Enterococcus faecalis* (44). This emphasises another aspect of AMR in *C. difficile*; its possible role as a reservoir of AMR genes for other pathogenic bacteria residing in the colon.

Previously, it has been reported that concordance between the presence of the *erm*(B) gene and an MLS_B_ resistance phenotype was low (45), likely due to the presence of multiple resistance mechanisms. In a previous study, however, the *erm*(B) gene was found in all *C. difficile* isolates with high-level resistance to both clindamycin and erythromycin, hinting that the gene may be associated with high-level MLS_B_ resistance (46). We also observed separation between *C. difficile* strains with high-level and low-level clindamycin and erythromycin resistance (**Figure 1**). Upon genomic analysis of a subset of strains, there was a strong correlation between the presence of an *erm*-class gene (*erm*(B), *erm*(G) and *erm*(52) genes) with high-level clindamycin resistance, which is usually accompanied by high-level erythromycin resistance, supporting the previous study (46). A macrolide efflux gene *mef*(G) was identified also in strains with high-level erythromycin resistance and not in erythromycin-susceptible strains, suggesting an association between the presence of this efflux protein and phenotypic erythromycin resistance, although the function of this gene was not characterised. Resistance determinants were not identified among strains with low-level clindamycin resistance, however, this underestimation is likely irrelevant, as the median clindamycin MIC in this population (8 mg/l) remained significantly lower than the clindamycin level in stools (approximately 240 mg/g of stool) (47). Besides MLS_B_, a separation between strains resistant and susceptible to rifaximin and fluoroquinolones was observed (**Figure 1**). The concordance between resistant phenotype and known genotype was also high, similar to a previous study (45).

Compared to the study at the same hospital in 2015, there was no difference in overall resistance prevalence (26), however, there was a slight increase in meropenem MICs and the emergence of carbapenem resistance. In a previous study, high-level imipenem resistance was associated with point substitutions in PBP1 and PBP3 (36). It appears that these substitutions only confer high-level resistance to imipenem and not to meropenem, although the linear regression analysis suggests that the Y721S substitution in PBP3 may have contributed to a slight increase in meropenem MIC in this study.

*C. difficile* remained susceptible to metronidazole, vancomycin and fidaxomicin, similar to the other parts of the world (48). Thus, these antimicrobials should remain effective treatments for CDI. There was a slight increase in vancomycin MIC reaching the clinical breakpoint, consistent with a previous study (26), however, this should have little impact on the treatment of CDI given that the faecal vancomycin concentration remains far greater than the MIC (>2,000 mg/l vs 2 mg/l) (49). The increase in vancomycin MIC in this study is in contrast to other hospitals in Thailand and this could reflect usage of vancomycin at the study site (22). Overuse of vancomycin can lead to the emergence of vancomycin-resistant *Enterococcus* spp., which can have a devastating effect on patients (50, 51). Therefore, vancomycin usage should be carefully monitored.

## Conclusion

A-B+CDT− *C. difficile* and NTCD remained prevalent in Thailand. Few binary toxin-positive strains (A+B+CDT+) were identified; one belonging to a known epidemic lineage and another a novel strain related to *C. difficile* RT 027. The most common RT in this study was *C. difficile* RT 017 (A-B+CDT-), a large proportion of which was resistant to MLS_B_, moxifloxacin and rifaximin. Many strains were also MDR. Such resistance may have played a role in the success of *C. difficile* RT 017 in Thailand. There was a strong concordance between the presence of *erm*-class genes and high-level clindamycin resistance, as well as significant concordance between point substitutions in gyrase subunits and RpoB with fluoroquinolone and rifaximin resistance, respectively. Resistance to antimicrobials suitable for the treatment of CDI was not detected.

## Acknowledgement

Parts of this study were performed using the facilities provided by the Pawsey Supercomputing Centre (Perth, Western Australia).

## Transparency declaration

T.V.R. has received grants from Cepheid; Merck; Otsuka; Roche; Sanofi and Summit for work outside that in this report. Other authors have no conflicts of interest to declare.

## References

1. Leffler DA, Lamont JT. 2015. *Clostridium difficile* Infection. N Engl J Med 373:287–8.

2. Braun V, Hundsberger T, Leukel P, Sauerborn M, von Eichel-Streiber C. 1996. Definition of the single integration site of the pathogenicity locus in *Clostridium difficile*. Gene 181:29–38.

3. Perelle S, Gibert M, Bourlioux P, Corthier G, Popoff MR. 1997. Production of a complete binary toxin (actin-specific ADP-ribosyltransferase) by *Clostridium difficile* CD196. Infect Immun 65:1402–7.

4. Kato N, Ou CY, Kato H, Bartley SL, Brown VK, Dowell VR, Jr., Ueno K. 1991. Identification of toxigenic *Clostridium difficile* by the polymerase chain reaction. J Clin Microbiol 29:33–7.

5. Stubbs S, Rupnik M, Gibert M, Brazier J, Duerden B, Popoff M. 2000. Production of actin-specific ADP-ribosyltransferase (binary toxin) by strains of *Clostridium difficile*. FEMS Microbiol Lett 186:307–12.

6. Imwattana K, Knight DR, Kullin B, Collins DA, Putsathit P, Kiratisin P, Riley TV. 2019. *Clostridium difficile* ribotype 017 - characterization, evolution and epidemiology of the dominant strain in Asia. Emerg Microbes Infect 8:796–807.

7. Kato H, Kato N, Watanabe K, Iwai N, Nakamura H, Yamamoto T, Suzuki K, Kim SM, Chong Y, Wasito EB. 1998. Identification of toxin A-negative, toxin B-positive *Clostridium difficile* by PCR. J Clin Microbiol 36:2178–82.

8. Lemee L, Dhalluin A, Testelin S, Mattrat M-A, Maillard K, Lemeland J-F, Pons J-L. 2004. Multiplex PCR targeting *tpi* (triose phosphate isomerase), *tcdA* (Toxin A), and *tcdB* (Toxin B) genes for toxigenic culture of *Clostridium difficile*. J Clin Microbiol 42:5710–4.

9. Stubbs SL, Brazier JS, O’Neill GL, Duerden BI. 1999. PCR targeted to the 16S-23S rRNA gene intergenic spacer region of *Clostridium difficile* and construction of a library consisting of 116 different PCR ribotypes. J Clin Microbiol 37:461–3.

10. Elliott B, Androga GO, Knight DR, Riley TV. 2017. *Clostridium difficile* infection: Evolution, phylogeny and molecular epidemiology. Infect Genet Evol 49:1–11.

11. Warny M, Pepin J, Fang A, Killgore G, Thompson A, Brazier J, Frost E, McDonald LC. 2005. Toxin production by an emerging strain of *Clostridium difficile* associated with outbreaks of severe disease in North America and Europe. Lancet 366:1079–84.

12. Bakker D, Corver J, Harmanus C, Goorhuis A, Keessen EC, Fawley WN, Wilcox MH, Kuijper EJ. 2010. Relatedness of human and animal *Clostridium difficile* PCR ribotype 078 isolates determined on the basis of multilocus variable-number tandem-repeat analysis and tetracycline resistance. J Clin Microbiol 48:3744–9.

13. Banawas SS. 2018. *Clostridium difficile* infections: a global overview of drug sensitivity and resistance mechanisms. Biomed Res Int 2018:8414257.

14. Thomas C, Stevenson M, Riley TV. 2003. Antibiotics and hospital-acquired *Clostridium difficile*-associated diarrhoea: a systematic review. J Antimicrob Chemother 51:1339–50.

15. Imwattana K, Knight DR, Kullin B, Collins DA, Putsathit P, Kiratisin P, Riley TV. 2020. Antimicrobial resistance in *Clostridium difficile* ribotype 017. Expert Rev Anti Infect Ther 18:17–25.

16. Slimings C, Riley TV. 2014. Antibiotics and hospital-acquired *Clostridium difficile* infection: update of systematic review and meta-analysis. J Antimicrob Chemother 69:881–91.

17. Centers for Disease Control and Prevention (CDC). 2019. Antibiotic resistance threats in the United States, 2019. U.S. Department of Health and Human Services, CDC, Atlanta, GA.

18. Cheng AC, Turnidge J, Collignon P, Looke D, Barton M, Gottlieb T. 2012. Control of fluoroquinolone resistance through successful regulation, Australia. Emerg Infect Dis 18:1453–1460.

19. Lew T, Putsathit P, Sohn KM, Wu Y, Ouchi K, Ishii Y, Tateda K, Riley TV, Collins DA. 2020. Antimicrobial susceptibilities of *Clostridium difficile* isolates from 12 Asia-Pacific countries in 2014 and 2015. Antimicrob Agents Chemother 64.

20. Putsathit P, Maneerattanaporn M, Piewngam P, Kiratisin P, Riley TV. 2017. Prevalence and molecular epidemiology of *Clostridium difficile* infection in Thailand. New Microbes New Infect 15:27–32.

21. Ngamskulrungroj P, Sanmee S, Putsathit P, Piewngam P, Elliott B, Riley TV, Kiratisin P. 2015. Molecular epidemiology of *Clostridium difficile* infection in a large teaching hospital in Thailand. PLoS One 10:e0127026.

22. Imwattana K, Wangroongsarb P, Riley TV. 2019. High prevalence and diversity of *tcdA*-negative and *tcdB*-positive, and non-toxigenic, *Clostridium difficile* in Thailand. Anaerobe 57:4–10.

23. Zellweger RM, Carrique-Mas J, Limmathurotsakul D, Day NPJ, Thwaites GE, Baker S, Southeast Asia Antimicrobial Resistance Network. 2017. A current perspective on antimicrobial resistance in Southeast Asia. J Antimicrob Chemother 72:2963–2972.

24. Putsathit P, Morgan J, Bradford D, Engelhardt N, Riley TV. 2015. Evaluation of the BD Max Cdiff assay for the detection of toxigenic *Clostridium difficile* in human stool specimens. Pathology 47:165–8.

25. Clinical and Laboratory Standards Institute (CLSI). 2012. Methods for Antimicrobial Susceptibility Testing of Anaerobic Bacteria, Approved Standard M11-A8, 8th ed. Clinical and Laboratory Standards Institute, Wayne, PA.

26. Putsathit P, Maneerattanaporn M, Piewngam P, Knight DR, Kiratisin P, Riley TV. 2017. Antimicrobial susceptibility of *Clostridium difficile* isolated in Thailand. Antimicrob Resist Infect Control 6:58.

27. O’Connor JR, Galang MA, Sambol SP, Hecht DW, Vedantam G, Gerding DN, Johnson S. 2008. Rifampin and rifaximin resistance in clinical isolates of *Clostridium difficile*. Antimicrob Agents Chemother 52:2813–7.

28. European Committee on Antimicrobial Susceptibility Testing. 16 May 2018. Clinical breakpoint tables, version 8.1. http://www.eucast.org/clinical_breakpoints/. Accessed 26 October 2018.

29. Goldstein EJC, Babakhani F, Citron DM. 2012. Antimicrobial activities of fidaxomicin. Clin Infect Dis 55 Suppl 2:S143–8.

30. Knight DR, Squire MM, Collins DA, Riley TV. 2016. Genome analysis of *Clostridium difficile* PCR ribotype 014 lineage in Australian pigs and humans reveals a diverse genetic repertoire and signatures of long-range interspecies transmission. Front Microbiol 7:2138.

31. Jain C, Rodriguez-R LM, Phillippy AM, Konstantinidis KT, Aluru S. 2018. High throughput ANI analysis of 90K prokaryotic genomes reveals clear species boundaries. Nat Commun 9.

32. Inouye M, Dashnow H, Raven LA, Schultz MB, Pope BJ, Tomita T, Zobel J, Holt KE. 2014. SRST2: Rapid genomic surveillance for public health and hospital microbiology labs. Genome Med 6:90.

33. Gupta SK, Padmanabhan BR, Diene SM, Lopez-Rojas R, Kempf M, Landraud L, Rolain JM. 2014. ARG-ANNOT, a new bioinformatic tool to discover antibiotic resistance genes in bacterial genomes. Antimicrob Agents Chemother 58:212–20.

34. Carver T, Harris SR, Berriman M, Parkhill J, McQuillan JA. 2012. Artemis: an integrated platform for visualization and analysis of high-throughput sequence-based experimental data. Bioinformatics 28:464–9.

35. Spigaglia P. 2016. Recent advances in the understanding of antibiotic resistance in *Clostridium difficile* infection. Ther Adv Infect Dis 3:23–42.

36. Isidro J, Santos A, Nunes A, Borges V, Silva C, Vieira L, Mendes AL, Serrano M, Henriques AO, Gomes JP, Oleastro M. 2018. Imipenem resistance in *Clostridium difficile* ribotype 017, Portugal. Emerg Infect Dis 24:741–745.

37. Roberts MC, Sutcliffe J, Courvalin P, Jensen LB, Rood J, Seppala H. 1999. Nomenclature for macrolide and macrolide-lincosamide-streptogramin B resistance determinants. Antimicrob Agents Chemother 43:2823–30.

38. Alcock BP, Raphenya AR, Lau TTY, Tsang KK, Bouchard M, Edalatmand A, Huynh W, Nguyen AV, Cheng AA, Liu S, Min SY, Miroshnichenko A, Tran HK, Werfalli RE, Nasir JA, Oloni M, Speicher DJ, Florescu A, Singh B, Faltyn M, Hernandez-Koutoucheva A, Sharma AN, Bordeleau E, Pawlowski AC, Zubyk HL, Dooley D, Griffiths E, Maguire F, Winsor GL, Beiko RG, Brinkman FSL, Hsiao WWL, Domselaar GV, McArthur AG. 2020. CARD 2020: antibiotic resistome surveillance with the comprehensive antibiotic resistance database. Nucleic Acids Res 48:D517–D525.

39. Valiente E, Cairns MD, Wren BW. 2014. The *Clostridium difficile* PCR ribotype 027 lineage: a pathogen on the move. Clin Microbiol Infect 20:396–404.

40. He M, Miyajima F, Roberts P, Ellison L, Pickard DJ, Martin MJ, Connor TR, Harris SR, Fairley D, Bamford KB, D’Arc S, Brazier J, Brown D, Coia JE, Douce G, Gerding D, Kim HJ, Koh TH, Kato H, Senoh M, Louie T, Michell S, Butt E, Peacock SJ, Brown NM, Riley T, Songer G, Wilcox M, Pirmohamed M, Kuijper E, Hawkey P, Wren BW, Dougan G, Parkhill J, Lawley TD. 2013. Emergence and global spread of epidemic healthcare-associated *Clostridium difficile*. Nat Genet 45:109–13.

41. Prakobsrikul N, Malathum K, Santanirand P, Chumnumwat S, Piebpien P, Montakantikul P. 2019. Correlation between antimicrobial consumption and the prevalence of carbapenem-resistant *Escherichia coli* and carbapenem-resistant *Klebsiella pneumoniae* at a university hospital in Thailand. J Clin Pharm Ther 44:292–299.

42. Knight DR, Imwattana K, Kullin B, Guerrero-Araya E, Paredes-Sabja D, Didelot X, Dingle KE, Eyre DW, Rodríguez C, Riley TV. 2020. The *Clostridioides difficile* species problem: global phylogenomic analysis uncovers three ancient, toxigenic, genomospecies. BioRxiv doi:https://doi.org/10.1101/2020.09.21.307223.

43. McDonald LC, Gerding DN, Johnson S, Bakken JS, Carroll KC, Coffin SE, Dubberke ER, Garey KW, Gould CV, Kelly C, Loo V, Shaklee Sammons J, Sandora TJ, Wilcox MH. 2018. Clinical practice guidelines for *Clostridium difficile* infection in adults and children: 2017 update by the Infectious Diseases Society of America (IDSA) and Society for Healthcare Epidemiology of America (SHEA). Clin Infect Dis 66:987–994.

44. Wasels F, Monot M, Spigaglia P, Barbanti F, Ma L, Bouchier C, Dupuy B, Mastrantonio P. 2014. Inter- and intraspecies transfer of a *Clostridium difficile* conjugative transposon conferring resistance to MLS_B_. Microb Drug Resist 20:555–60.

45. Knight DR, Kullin B, Androga GO, Barbut F, Eckert C, Johnson S, Spigaglia P, Tateda K, Tsai PJ, Riley TV. 2019. Evolutionary and genomic insights into *Clostridioides difficile* sequence type 11: a diverse zoonotic and antimicrobial-resistant lineage of global one health importance. mBio 10.

46. Solomon K, Fanning S, McDermott S, Murray S, Scott L, Martin A, Skally M, Burns K, Kuijper E, Fitzpatrick F, Fenelon L, Kyne L. 2011. PCR ribotype prevalence and molecular basis of macrolide-lincosamide-streptogramin B (MLS_B_) and fluoroquinolone resistance in Irish clinical *Clostridium difficile* isolates. J Antimicrob Chemother 66:1976–82.

47. Kager L, Liljeqvist L, Malmborg AS, Nord CE. 1981. Effect of clindamycin prophylaxis on the colonic microflora in patients undergoing colorectal surgery. Antimicrob Agents Chemother 20:736–40.

48. Freeman J, Vernon J, Pilling S, Morris K, Nicholson S, Shearman S, Longshaw C, Wilcox MH, Pan-European Longitudinal Surveillance of Antibiotic Resistance among Prevalent Clostridium difficile Ribotypes Study G. 2018. The ClosER study: results from a three-year pan-European longitudinal surveillance of antibiotic resistance among prevalent *Clostridium difficile* ribotypes, 2011-2014. Clin Microbiol Infect 24:724–731.

49. Gonzales M, Pepin J, Frost EH, Carrier JC, Sirard S, Fortier LC, Valiquette L. 2010. Faecal pharmacokinetics of orally administered vancomycin in patients with suspected *Clostridium difficile* infection. BMC Infect Dis 10:363.

50. Chotiprasitsakul D, Santanirand P, Thitichai P, Rotjanapan P, Watcharananan S, Siriarayapon P, Chaihongsa N, Sirichot S, Chitasombat M, Chantharit P, Malathum K. 2016. Epidemiology and control of the first reported vancomycin-resistant *Enterococcus* outbreak at a tertiary-care hospital in Bangkok, Thailand. Southeast Asian J Trop Med Public Health 47:494–502.

51. Hemapanpairoa J, Changpradub D, Thunyaharn S, Santimaleeworagun W. 2019. Vancomycin-resistant enterococcal infection in a Thai university hospital: clinical characteristics, treatment outcomes, and synergistic effect. Infect Drug Resist 12:2049–2057.

