## Supplementary for "Molecular characterisation of, and antimicrobial resistance in, *Clostridioides difficile* from Thailand, 2017-2018"

**Table S1 – Summary of genome metrics.**

| Strain ID | Accession | Collection year | Toxin profile | ST | Clade | Genome size (Mbp) | $N_{50}$ (bp) | N of contigs | GC% |
|---|---|---|---|---|---|---|---|---|---|
| MAR006 | ERS5247348 | 2017 | A-B+CDT- | 37 | 4 | 4.32 | 314,706 | 36 | 28.93 |
| MAR014 | ERS5247349 | 2017 | A-B+CDT- | 37 | 4 | 4.46 | 314,836 | 37 | 28.63 |
| MAR024 | ERS5247350 | 2017 | A-B+CDT- | 37 | 4 | 4.21 | 295,420 | 36 | 28.53 |
| MAR027 | ERS5247351 | 2017 | A-B+CDT- | 37 | 4 | 4.38 | 283,659 | 37 | 28.49 |
| MAR037 | ERS5247352 | 2017 | A-B+CDT- | 37 | 4 | 4.49 | 219,221 | 43 | 28.87 |
| MAR040 | ERS5247353 | 2017 | A-B+CDT- | 37 | 4 | 4.22 | 294,666 | 39 | 28.54 |
| MAR063 | ERS5247354 | 2017 | A-B+CDT- | 37 | 4 | 4.15 | 219,220 | 37 | 28.39 |
| MAR064 | ERS5247355 | 2017 | A-B+CDT- | 37 | 4 | 4.16 | 309,809 | 32 | 28.50 |
| MAR065 | ERS5247356 | 2017 | A-B+CDT- | 37 | 4 | 4.49 | 217,096 | 55 | 28.60 |
| MAR076 | ERS5247357 | 2017 | A-B+CDT- | 37 | 4 | 4.23 | 183,348 | 39 | 28.31 |
| MAR081 | ERS5247358 | 2017 | A-B+CDT- | 37 | 4 | 4.21 | 243,481 | 35 | 28.52 |
| MAR084 | ERS5247359 | 2017 | A-B+CDT- | 37 | 4 | 4.29 | 215,624 | 47 | 28.66 |
| MAR099 | ERS5247360 | 2017 | A-B+CDT- | 37 | 4 | 4.24 | 376,582 | 31 | 28.83 |
| MAR138 | ERS5247361 | 2017 | A-B+CDT- | 37 | 4 | 4.32 | 219,010 | 39 | 28.68 |
| MAR140 | ERS5247362 | 2017 | A-B+CDT- | 37 | 4 | 4.36 | 202,573 | 43 | 28.71 |
| MAR153 | ERS5247363 | 2017 | A-B+CDT- | 37 | 4 | 4.28 | 244,797 | 34 | 28.77 |
| MAR169 | ERS5247364 | 2017 | A-B+CDT- | 37 | 4 | 4.28 | 302,420 | 33 | 28.78 |
| MAR178 | ERS5247365 | 2018 | A-B+CDT- | 37 | 4 | 4.28 | 302,420 | 28 | 28.78 |
| MAR191 | ERS5247366 | 2018 | A-B+CDT- | 37 | 4 | 4.18 | 210,823 | 36 | 28.58 |
| MAR196 | ERS5247367 | 2018 | A-B+CDT- | 37 | 4 | 4.43 | 314,700 | 33 | 28.62 |
| MAR199 | ERS5247368 | 2018 | A-B+CDT- | 37 | 4 | 4.28 | 244,647 | 36 | 28.77 |
| MAR213 | ERS5247369 | 2018 | A+B+CDT+ | 692 | 1* | 4.07 | 184,956 | 54 | 28.29 |
| MAR214 | ERS5247370 | 2018 | A+B+CDT- | 2 | 1 | 4.28 | 2,281,307 | 30 | 28.74 |
| MAR217 | ERS5247371 | 2018 | A-B+CDT- | 37 | 4 | 4.32 | 194,208 | 51 | 28.52 |
| MAR218 | ERS5247372 | 2018 | A-B+CDT- | 37 | 4 | 4.32 | 217,855 | 47 | 28.52 |
| MAR225 | ERS5247373 | 2018 | A-B+CDT- | 37 | 4 | 4.29 | 293,537 | 36 | 28.64 |
| MAR235 | ERS5247374 | 2018 | A-B+CDT- | 37 | 4 | 4.29 | 244,785 | 34 | 28.63 |
| MAR245 | ERS5247375 | 2018 | A-B+CDT- | 37 | 4 | 4.48 | 183,229 | 46 | 28.61 |
| MAR251 | ERS5247376 | 2018 | A-B+CDT- | 37 | 4 | 4.35 | 221,550 | 46 | 28.68 |
| MAR272 | ERS5247377 | 2018 | A-B+CDT- | 37 | 4 | 4.30 | 283,687 | 38 | 28.51 |
| MAR273 | ERS5247378 | 2018 | A-B+CDT- | 37 | 4 | 4.23 | 535,521 | 30 | 28.29 |
| MAR286 | ERS5247379 | 2018 | A-B+CDT- | 37 | 4 | 4.17 | 285,837 | 38 | 28.56 |

Note: * Clade 1 by MLST scheme.

**Table S1 (continued) – Summary of genome metrics.**

| Strain ID | Accession | Collection year | Toxin profile | ST | Clade | Genome size (Mbp) | $N_{50}$ (bp) | N of contigs | GC% |
|---|---|---|---|---|---|---|---|---|---|
| MAR297 | ERS5247380 | 2018 | A-B+CDT- | 37 | 4 | 4.35 | 293,509 | 40 | 28.68 |
| MAR300 | ERS5247381 | 2018 | A-B+CDT- | 37 | 4 | 4.32 | 211,511 | 38 | 28.69 |
| MAR302 | ERS5247382 | 2018 | A-B+CDT- | 37 | 4 | 4.17 | 376,582 | 33 | 28.60 |
| MAR306 | ERS5247383 | 2018 | A-B+CDT- | 37 | 4 | 4.49 | 165,297 | 64 | 28.74 |
| MAR310 | ERS5247384 | 2018 | A-B+CDT- | 37 | 4 | 4.20 | 352,120 | 32 | 28.56 |
| **Median** | | | | | | 4.29 | 244,797 | 37 | 28.62 |
| **Range** | | | | | | 4.07 – 4.49 | 165,297 – 2,281,307 | 28 – 64 | 28.29 – 28.93 |

**Table S2 – Comparison of antimicrobial resistance phenotype and genotype among sequenced *C. difficile* strains.**

| Strain ID | Accession | $MLS_B$ | | | MOX | | RIF | | MER | |
|---|---|---|---|---|---|---|---|---|---|---|
| | | MIC (CLI) | MIC (ERY) | Genotype | MIC | Genotype[1] | MIC | Genotype[2] | MIC | Genotype[3] |
| MAR006 | ERS5247348 | >256 | >256 | *erm*(B) | 32 | T82I(A) | >64 | H502N, R505K | 4 | A555T(1) |
| MAR014 | ERS5247349 | >256 | >256 | *erm*(B) | 32 | T82I(A) | >64 | H502N, R505K | 4 | A555T(1) |
| MAR024 | ERS5247350 | 256 | >256 | *erm*(B) | 32 | T82I(A) | >64 | H502N, R505K | 8 | - |
| MAR027 | ERS5247351 | 256 | >256 | *erm*(B) | 2 | - | 0.03 | - | 4 | - |
| MAR037 | ERS5247352 | >256 | >256 | *erm*(B) | 32 | T82I(A) | >64 | H502N, R505K | 4 | A555T(1) |
| MAR040 | ERS5247353 | 256 | >256 | *erm*(B) | 32 | T82I(A) | >64 | H502N, R505K | 8 | - |
| MAR063 | ERS5247354 | 8 | 1 | - | 2 | - | 0.015 | - | 4 | - |
| MAR064 | ERS5247355 | 128 | >256 | *erm*(B) | 8 | - | >64 | H502N, R505K | 4 | - |
| MAR065 | ERS5247356 | >256 | >256 | *erm*(B) | 16 | T82I(A) | >64 | H502N, R505K | 4 | A555T(1) |
| MAR076 | ERS5247357 | 16 | 2 | - | 2 | - | 0.008 | - | 4 | - |
| MAR081 | ERS5247358 | >256 | >256 | *erm*(B) | 2 | - | 0.008 | - | 4 | - |
| MAR084 | ERS5247359 | >256 | >256 | *erm*(B) | 32 | T82I(A) | >64 | H502N, R505K | 4 | - |
| MAR099 | ERS5247360 | 256 | >256 | *erm*(G), *msr*(D) | 2 | - | 0.008 | - | 4 | - |
| MAR138 | ERS5247361 | >256 | >256 | *erm*(B) | 32 | T82I(A) | >64 | H502N, R505K | 4 | A555T(1) |
| MAR140 | ERS5247362 | >256 | >256 | *erm*(B) | 32 | T82I(A) | >64 | H502N, R505K | 4 | A555T(1) |
| MAR153 | ERS5247363 | >256 | >256 | *erm*(G), *msr*(D) | 32 | T82I(A) | >64 | H502N, R505K | 8 | Y721S(3) |
| MAR169 | ERS5247364 | >256 | >256 | *erm*(G), *msr*(D) | 32 | T82I(A) | >64 | H502N, R505K | 8 | Y721S(3) |
| MAR178 | ERS5247365 | >256 | >256 | *erm*(G), *msr*(D) | 32 | T82I(A) | >64 | H502N, R505K | 8 | Y721S(3) |

Note: [1] point substitutions on either GyrA (A) or GyrB (B) subunit; [2] point substitutions on RpoB; [3] point substitutions on either PBP1 (1) or PBP3 (3); $MLS_B$, macrolide-lincosamide-streptogramin B; CLI, clindamycin; ERY, erythromycin; MOX, moxifloxacin; MER, meropenem.

**Table S2 (continued) – Comparison of antimicrobial resistance phenotype and genotype among sequenced *C. difficile* strains.**

| Strain ID | Accession | $MLS_B$ | | | MOX | | RIF | | MER | |
|---|---|---|---|---|---|---|---|---|---|---|
| | | MIC (CLI) | MIC (ERY) | Genotype | MIC | Genotype[1] | MIC | Genotype[2] | MIC | Genotype[3] |
| MAR191 | ERS5247366 | 256 | >256 | *erm*(B) | 2 | - | 0.03 | - | 4 | - |
| MAR196 | ERS5247367 | >256 | >256 | *erm*(B) | 32 | T82I(A) | >64 | H502N, R505K | 4 | A555T(1) |
| MAR199 | ERS5247368 | >256 | >256 | *erm*(G), *msr*(D) | 32 | T82I(A) | >64 | H502N, R505K | 8 | Y721S(3) |
| MAR213 | ERS5247369 | 8 | 2 | - | 2 | - | 0.03 | - | 4 | - |
| MAR214 | ERS5247370 | >256 | 0.5 | *erm*(B) | 16 | D426N(B) | 0.03 | - | 2 | - |
| MAR217 | ERS5247371 | >256 | >256 | *erm*(B) | 2 | - | 0.03 | - | 8 | - |
| MAR218 | ERS5247372 | >256 | >256 | *erm*(B) | 2 | - | 0.03 | - | 8 | - |
| MAR225 | ERS5247373 | >256 | >256 | *erm*(52) | 32 | T82I(A) | >64 | H502N, R505K | 8 | Y721S(3) |
| MAR235 | ERS5247374 | >256 | 64 | *erm*(52) | 32 | T82I(A) | >64 | H502N, R505K | 8 | Y721S(3) |
| MAR245 | ERS5247375 | 256 | >256 | *erm*(B) | 32 | T82I(A) | >64 | H502N, R505K | 4 | A555T(1) |
| MAR251 | ERS5247376 | 256 | >256 | *erm*(B) | 32 | T82I(A) | >64 | H502N, R505K | 4 | A555T(1) |
| MAR272 | ERS5247377 | 8 | >256 | *mef*(G) | 2 | - | 0.015 | - | 4 | - |
| MAR273 | ERS5247378 | 8 | 1 | - | 2 | - | 0.015 | - | 4 | - |
| MAR286 | ERS5247379 | >256 | >256 | *erm*(B) | 2 | - | 0.03 | - | 4 | - |
| MAR297 | ERS5247380 | 256 | >256 | *erm*(B) | 32 | T82I(A) | >64 | H502N, R505K | 4 | A555T(1) |
| MAR300 | ERS5247381 | >256 | >256 | *erm*(B) | 32 | T82I(A) | >64 | H502N, R505K | 4 | A555T(1) |
| MAR302 | ERS5247382 | >256 | 64 | *erm*(52) | 32 | T82I(A) | >64 | H502N, R505K | 16 | Y721S(3) |
| MAR306 | ERS5247383 | >256 | >256 | *erm*(B) | 32 | T82I(A) | >64 | H502N, R505K | 4 | A555T(1) |
| MAR310 | ERS5247384 | 16 | >256 | *mef*(G) | 2 | - | 0.03 | - | 4 | - |

Note: [1] point substitutions on either GyrA (A) or GyrB (B) subunit; [2] point substitutions on RpoB; [3] point substitutions on either PBP1 (1) or PBP3 (3); $MLS_B$, macrolide-lincosamide-streptogramin B; CLI, clindamycin; ERY, erythromycin; MOX, moxifloxacin; MER, meropenem.
